# Widespread translational remodeling during human neuronal differentiation

**DOI:** 10.1101/156190

**Authors:** John D. Blair, Dirk Hockemeyer, Jennifer A. Doudna, Helen S. Bateup, Stephen N. Floor

## Abstract

Faithful cellular differentiation requires precise coordination of changes in gene expression. However, the relative contributions of transcriptional and translational regulation during human cellular differentiation are unclear. Here, we induced forebrain neuronal differentiation of human embryonic stem cells (hESCs) and characterized genomewide RNA and translation levels during neurogenesis. We find that thousands of genes change at the translation level across differentiation without a corresponding change in RNA level. Specifically, we identify mTOR complex 1 signaling as a key driver for elevated translation of translation-related genes in hESCs. In contrast, translational repression in active neurons is mediated by transcript 3′ UTRs, through regulatory sequences. Together, our findings identify a functional role for the dramatic 3′ UTR extensions that occur during brain development, and provide insights to interpret genetic variants in post-transcriptional control factors that influence neurodevelopmental disorders and diseases.

## Introduction

Specification of differentiated cell types requires tightly coordinated execution of gene expression programs. The balance and timing of proliferation and differentiation must be exquisitely controlled to promote proper development and avoid unrestricted growth. Coordination of gene expression is especially important in the development and maintenance of post-mitotic cells of neuronal lineages, as deviant proteostasis can lead to diverse pathologies including autism and neurodegenerative disease (Darnell and Klann, 2013; Freibaum and Taylor, 2017; Gkogkas et al., 2013; Kelleher and Bear, 2008). To achieve this coordination, cellular systems regulate each step of gene expression, including transcription, splicing and transcript processing, translation, and mRNA or protein degradation.

The brain contains the most complex mRNA repertoire of any human tissue (Ramskold et al., 2009). Transcript 3′ untranslated regions (UTRs) are lengthened considerably in the brain and during neuronal differentiation, where individual 3′ UTRs can exceed 20 kb (Hilgers et al., 2011; Miura et al., 2013). Extension of 3′ UTRs is driven by changes in expression levels of RNA-binding proteins that influence polyadenylation site selection or selective degradation of elongated 3′ UTRs (Hilgers et al., 2012; Mayr, 2016; Tian and Manley, 2017). Transcript 3′ UTRs regulate translation level, RNA degradation, RNA localization, and protein complex formation, therefore this expansion in 3′ UTR size also increases the post-transcriptional regulatory potential of the brain (Mayr, 2016; Miura et al., 2014; Tian and Manley, 2017). Defective 3′ UTR formation leads to nervous system dysfunction (Van Epps et al., 2010) and widespread shortening of 3′ UTRs is observed upon a shift to proliferation in diverse contexts including cancers and other diseases (Elkon et al., 2013; Gennarino et al., 2015; Mayr and Bartel, 2009; Sandberg et al., 2008; Singh et al., 2009).

Despite the importance of brain-specific 3′ UTRs, how the expanded regulatory potential of mRNA 3′ UTRs influences gene expression during neuronal differentiation remains incompletely understood. Transcript 3′ UTRs have been demonstrated to influence subcellular localization of transcripts in neurons (An et al., 2008; Taliaferro et al., 2016). However, 3′ UTRs can also affect protein translation or mRNA degradation by recruiting RNA-binding proteins and miRNAs (Floor and Doudna, 2016; Mayr, 2016; Mayr and Bartel, 2009; Sandberg et al., 2008; Shenoy and Blelloch, 2014). It is unknown how brain-specific 3′ UTRs affect these processes, in part because RNA-sequencing analysis measures steady-state transcript levels but not their translation level. Furthermore, ribosome profiling, which measures the location of ribosomes on mRNAs using RNase-protected footprints, cannot measure translational effects of 3′ UTRs as they are digested. Therefore, the connection between brain-specific 3′ UTR extension and protein translation is unclear.

Here, we systematically quantified transcriptional and translational changes during human forebrain neuronal differentiation by performing parallel RNA-seq, ribosome profiling, and polysome profiling with transcript isoform resolution (TrIP-seq; (Floor and Doudna, 2016). We find that the mechanistic target of rapamycin compex 1 (mTORC1) signaling pathway drives high-level translation of translation-related genes selectively in human embryonic stem cells (hESCs), while transcriptional and translational changes are correlated for many other genes. At the transcript level, the presence of long 5′ UTRs repress translation in pluripotent and differentiated cells. In contrast, 3′ UTRs exert the strongest effect of any feature tested on translation in synaptically-active neural cultures, but have a minimal effect in hESCs. Cell-type-selective translational repression by 3′ UTRs is driven by an increase in the density of binding sites for regulatory RNA-binding proteins and miRNAs as opposed to 3′ UTR length. Therefore, the augmented post-transcriptional regulatory potential of brain-specific 3′ UTR extensions downregulates protein production, which may be crucial for tightly regulating proteostasis in highly specialized, post-mitotic cells such as neurons.

## Methods

### hESC cell culture

Human embryonic stem cell culture was carried out as previously described (Blair et al., 2016) in WIBR3 hESCs (NIH stem cell registry # 0079; (Lengner et al., 2010). Briefly, all hESC lines were maintained on a layer of inactivated mouse embryonic fibroblasts (MEFs) in hESC medium composed of DMEM/F12 supplemented with 20% KnockOut Serum Replacement, 2 mM glutamine, 1% non-essential amino acids, 0.1 mM β-mercaptoethanol, 1000 U/ml penicillin/streptomycin, and 4 ng/ml FGF2. Cultures were passaged every 7 days with collagenase type IV and gravitational sedimentation by washing 3 times in wash media composed of DMEM/F12 supplemented with 5% fetal bovine serum and 1000 U/ml penicillin/streptomycin. hESCs were regularly tested for mycoplasma contamination.

### Neuronal differentiation and culture

Neural induction was performed as described previously (Chambers et al., 2011), with minor alterations. Single hESCs were initially plated at a density of 50,000/cm2 (1.9x105 cells/well of a 12-well plate) and maintained in complete conditioned hESC media until >90% confluent. hESCs were transferred to induction media supplemented with 100 ng/μl Noggin and 10 μM SB431542 with daily media changes for 10 days. The composition of induction media changes throughout induction with 100% induction media A (A) from days 1-4, 75% A and 25% induction media B (B) on days 5-6, 50% A and 50% B on days 7-8 and 25% A and 75% B on days 9-10. Induction media A is composed of Knockout DMEM with 15% KSR, 2 mM l-glutamine, 1% non-essential amino acids, 1000 U/ml penicillin/streptomycin and 55 μM β-mercaptoethanol. Induction media B is composed of 50% DMEM/F12 media, 50% Neurobasal media 1x N-2 Supplement, 1x Glutamax, 1000 U/ml penicillin/streptomycin, 0.2% human Insulin and 0.075% bovine serum albumin as previously described (Chambers et al., 2011). After neural induction was complete, cells were dissociated with Accutase, spun down for 4 minutes at 800 rpm, resuspended in induction media B supplemented with 25 ng/ml FGF2 and 40 ng/ml EGF and replated on matrigel at 1:2. Two-thirds of media supplemented with FGF2 and EGF was changed every other day. Cells were passaged as such every 5 days until passage 4, when they were split at 1:3.

Neuronal differentiation from NPCs commenced at passage 8. NPCs were plated at low density – 80,000/cm2 (8x105 cells/well of a 6- well plate) in 4 ml of growth media. For imaging, cells were plated at the same density on a 12mm glass coverslip in a 24-well plate. Growth media (N2B27 media) is composed of 50% DMEM/F12, 50% Neurobasal Media, 1x N-2 Supplement, 1x B-27 Supplement, 1x Glutamax, 1000 U/ml penicillin/streptomycin and 0.075% BSA w/v. For the first 12 days, growth media was supplemented with 20 ng/ml BDNF and 20 ng/ml NT-3. For subsequent media changes, N2B27 media was used with no additional growth factors. All neuronal media changes occurred every 4 days with a half-volume change.

### Replicates

For all experiments both technical and biological replicates were used. Technical replicates consisted of cells that were plated at the same time treated with identical experimental handling and conditions. Biological replicates were cells plated one passage apart with matched cell numbers and experimental handling.

### Electrophysiology in day 50 neural cultures

Whole cell patch clamp recordings were made at room temperature from visually identified neurons grown on coverslips for 54-56 days after differentiation from NPCs. Cells were superfused with ACSF containing (in mM): 123 NaCl, 25 D-Glucose, 10 HEPES, 25 NaHCO3, 5 KCl, 1 Na2H2PO4, 2 CaCl2, and 1 MgCl2. Internal solution contained (in mM): 135 KMeSO4, 10 HEPES, 4 MgCl2, 4 Na-ATP, 0.4 Na-GTP, 10 phosphocreatine-Na2, and 1 EGTA. For current clamp recordings, the membrane potential was adjusted to -70mV and one second steps of depolarizing current (from +5-50pA) were injected to elicit action potentials. To measure excitatory synaptic activity, voltage clamp recordings were obtained at -70mV. Spontaneous excitatory synaptic currents were recorded before and five minutes after wash-in of the AMPA receptor blocker NBQX (10uM).

### Western blotting

Cells were harvested in lysis buffer containing 2 mM EDTA, 2 mM EGTA, 1% Triton-X, and 0.5% SDS in PBS with Halt phosphatase inhibitor cocktail and Complete mini EDTA-free protease inhibitor cocktail. Total protein was determined by BCA assay, and 10 μg of protein in Laemmli sample buffer were loaded onto 4–15% Criterion TGX gels. Proteins were transferred to PVDF membranes, blocked in 5% milk in TBS-Tween for one hour at room temperature (RT), and incubated with primary antibodies diluted in 5% milk in TBS-Tween overnight at 4°C. The following day, membranes were incubated with HRP-conjugated secondary antibodies for one hour at RT, washed, incubated with chemiluminesence substrate and developed on GE Amersham Hyperfilm ECL. Membranes were stripped with 6M guanidine hydrochloride to re-blot on subsequent days.

### Harvesting cells

Cells were treated with 100 ug/ml cycloheximide dissolved in pre-warmed media at 37 °C for 5 minutes before harvesting and 100 ug/ml cycloheximide was added to all downstream buffers. hESCs were dissociated from the feeder layer using 1 mg/ml Collagenase Type IV for 20 minutes followed by gravity sedimentation 3 times in wash media composed of DMEM/F12 supplemented with 5% fetal bovine serum and 1000 U/ml penicillin/streptomycin. NPCs and neural cultures were dissociated with Accutase and spun down for 4 minutes at 800 rpm. Cell pellets from ~10 million cells were washed with PBS, spun down and resuspended in 500 ul hypotonic lysis buffer on ice (10 mM HEPES pH 7.9, 1.5 mM MgCl2, 10 mM KCl, 0.5 mM DTT, 1% Triton X-100, 100 ug/ml cycloheximide). All samples were then incubated on ice for 10 minutes, triturated ten times through a 26G needle, spun at 1500g for 5 minutes at 4 °C (to pellet nuclei and cell bodies), and the supernatant was transferred to a new tube, which was used for either polysome or ribosome profiling in parallel. 20-30 ul of the cytoplasmic lysate was added to 300 ul Trizol reagent, and the nuclei were resuspended in 300 ul Trizol reagent, to generate cytoplasmic and nuclear RNA samples, respectively, and RNA was purified using the manufacturer’s protocol.

### Polysome profiling

Polysomes were separated using sucrose gradient centrifugation as previously described (Floor and Doudna, 2016). Sucrose gradients from 10-50% were made in 100 mM KCl, 20 mM HEPES pH 7.6 and 5 mM MgCl2 with 100 μg/ml cycloheximide and 0.66 U/μl Superase:In (Thermo-Fisher) and pre-chilled in centrifuge buckets for at least 30 minutes before use. 100 μl of lysate was then applied to the top of a 12 ml 10-50% sucrose gradient. Tubes were spun for 2 hours at 36,000 RPM (221,632 g) in a SW-41 rotor. The bottom of the tube was punctured and 2M sucrose pumped in using a peristaltic pump. Absorbance at 260 nm was monitored using a Brandel (Gaithersburg, MD) gradient fractionator and ISCO (Lincoln, NE) UA-6 detector. Peaks were fractionated into separate tubes, which were ethanol precipitated by adding 2 μl glycoblue (Thermo), 1:10 volumes 3M NaOAc pH 5.2 and 2 volumes 100% ethanol. Samples were resuspended, DNase treated using RQ1 RNase-free DNase (Promega), acid phenol: chloroform extracted, and ethanol precipitated as above. Samples were then resuspended in 20 μl DEPC-treated water and RNA concentration was measured using a Qubit (Life Technologies). Equal volumes of samples from the 2-4 ribosomes and the 5-8+ ribosomes fractions were then pooled.

### RNA-seq library prep

An Agilent Bioanalyzer was used to determine the RNA integrity (RIN) for each sample, which was typically > 6. Libraries were prepared using the TruSeq Stranded Total RNA Library Prep Kit (Illumina) starting with 100 ng of RNA from each sample or pool with 6 minutes fragmentation time and 13 PCR cycles.

### Ribosome profiling library prep

Ribosome profiling libraries were prepared using a standard protocol (McGlincy and Ingolia, 2017), with minor modifications. Sucrose gradient purification of 80S monosomes was used instead of a sucrose cushion and RNA was extracted from the 80S peak as above for polysome-derived material. A larger fragment size was excised from the initial ribosome protected RNA gel, from about 20nt to 40 nt, as reports have shown some variability in the protected footprint size (Guydosh and Green, 2014; Lareau et al., 2014). Following ligation of a linker containing a 5 N unique molecular identifier (UMI), excess adapter was digested by adding yeast 5′ deadenylase (Epicenter) and RecJ (Epicenter) and the material was purified using a Zymo Oligo Clean & Concentrator column. Ribosomal RNA depletion using RiboZero (Epicenter) was performed on each sample before reverse transcription. Matched RNA-seq libraries were constructed with the ribosome profiling library prep using RNA that was fragmented by incubating for 20 minutes at 95 degrees in 1 mM EDTA, 6 mM Na2CO3, 44 mM NaHCO3, pH 9.3. PCR was performed for 8 or 10 PCR cycles, depending on the library. The RNase digestion repeatedly overdigested ribosomes derived from 50 day old neural cultures causing them to fall apart during purification, precluding collection of ribosome protected footprints from these cells. Metagene plots were made using metagenemaker (https://github.com/stephenfloor/metagene-maker).

### High-throughput sequencing

All libraries were sequenced on an Illumina HiSeq 4000. Index bleed issues were corrected for in ribosome profiling libraries using read-internal indexes. Index bleed appeared to be a minor issue in polysomal TrIP-seq data, as, for example, OCT4 expression was tightly restricted to hESC libraries (Figure 1C). Total RNA libraries were sequenced in 100bp paired-end format, while ribosome profiling libraries were sequenced in 50bp single-read format. When necessary, multiple sequencing runs were performed and reads for each library were merged before downstream analysis. Sequencing was performed at the Vincent J. Coates Genomics Sequencing Laboratory at UC Berkeley. All sequencing reads are available through the NCBI GEO with accession number GSE100007.

**Figure 1:**
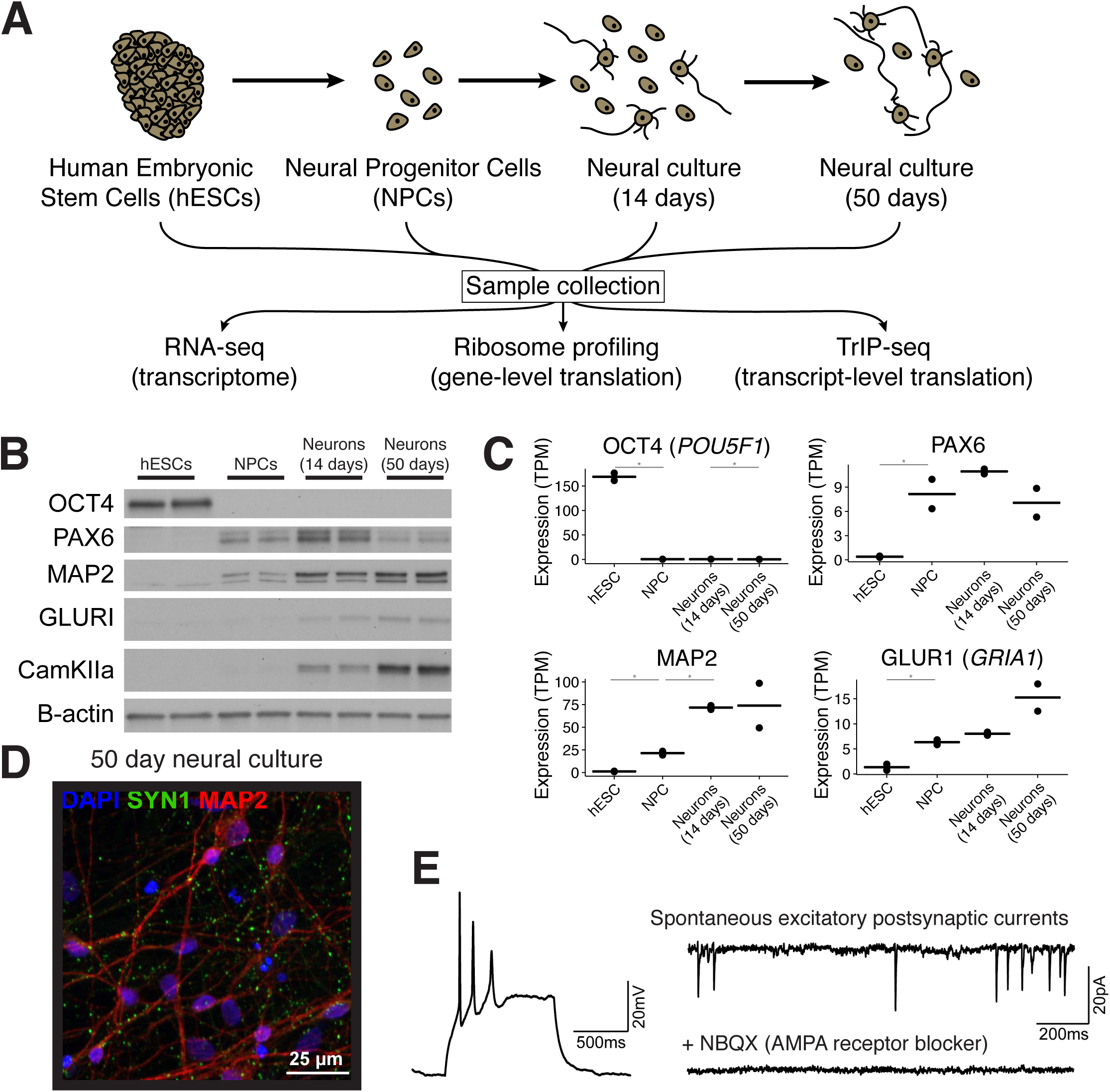
Measuring global changes in transcription and translation during human neuronal differentiation. (A) The experimental design. Human embryonic stem cells (hESCs) were differentiated into neural progenitor cells using dual-SMAD inhibition followed by neural induction. Samples were collected from each cell population and RNA-seq, ribosome profiling, and TrIP-seq libraries were prepared from each. (B) Western blotting of five cell-type markers during differentiation. (C) Examples of four marker genes from cytoplasmic RNA-seq. TPM: gene-level expression in transcripts-per-million. *: identified as differentially expressed by DESeq2 at p < 0.01. Bar: mean expression; points: expression in each replicate. (D) A representative image of 50-day old in vitro differentiated human neural cultures. Green: synapsin; red: MAP2; blue: DAPI. (E) Left, example whole cell current clamp recording of a neuron showing action potentials elicited by a 25pA depolarizing current injection. Right, example spontaneous excitatory post-synaptic currents (sEPSCs) showing network connectivity. sEPSCs were abolished by blocking AMPA receptors with NBQX (bottom right).See also Figure S1.

### Ribosome profiling data processing and analysis

Adapters were removed with Cutadapt 1.13 (Martin, 2011) (--trim-n -u 3 --minimum-length 15), retaining the UMI. PCR duplicate reads were removed using PRINSEQ (Schmieder and Edwards, 2011) (-derep 1 - no_qual_header). Five-base UMIs were then removed with Cutadapt (-- trim-n -u -5 --minimum-length 15). Trimmed reads that aligned to human rRNA and then to RMSK-derived regions were removed using Bowtie2 v 2.2.6 (Langmead and Salzberg, 2012) (-L 10 -i S,1,0.5). Remaining reads were quantified using Kallisto 0.43 (Bray et al., 2016) (-bias --fr-stranded --single -l <avg> -s <stdev>) using indexes made from all Ensembl GRCh38 v84 transcripts for both ORF and 5′ UTR regions. Regions were merged if they differed by less than 100 bp for ORFs and 50 bp for 5′ UTRs prior to index generation. Quantified transcript regions were aggregated to the gene level using tximport and analyzed using DESeq2 (Love et al., 2014). Statistically significant changes were called at p < 0.01. Biological duplicate samples were highly correlated (Figure S1D). The 5′ UTR to ORF ratio was computed by dividing 5′ UTR-quantified read counts per gene by ORF-quantified read counts per gene. Reads were aligned to the genome for visualization using HISAT2 v2.0.4 (Kim et al., 2015) and visualized with IGV (Thorvaldsdottir et al., 2013). Bedtools 2.21 (Quinlan, 2014) and Samtools 0.1.19 and 1.3 (Li et al., 2009) were also used.

### TrIP-seq polysome data processing and analysis

Paired-end adapters were removed with Cutadapt 1.13 (--minimum-length 50), and trimmed reads that aligned to human rRNA or to RMSK-derived regions were removed using Bowtie2 v 2.2.6. Remaining reads were quantified using Kallisto 0.42 (--bias) using indexes made from all Ensembl GRCh38 v84 transcripts. Quantified transcript regions were clustered (next section) or analyzed using DESeq2, or aggregated to the gene level using tximport and analyzed using DESeq2. Statistically significant changes were called at p < 0.01. Biological duplicate samples were highly correlated (Figure S1E). Reads were aligned to the genome for visualization using HISAT2 v2.0.4 and visualized with IGV. Classes of genes representing different cell types (Figure S1C) were identified in (Pollen et al., 2014) or (Mallon et al., 2013). The median expression within each gene class is plotted. Python programs used in this analysis can be downloaded at http://github.com/stephenfloor/tripseq-analysis.

### Hierarchical clustering and analysis

Clustering was performed essentially as described (Floor and Doudna, 2016), using Euclidean distance. Transcripts were clustered across the monosomal, low polysomal, and high polysomal samples to reflect transcript abundance across the polysome profile. The mean-subtracted, regularized-log transformation (rlog; DESeq2) of each transcript was clustered. Cluster number was determined by merging clusters below a specified dendrogram height until mergers combined clusters with different average profiles. Transcripts from clusters predominantly mapping to monosome, polysome-low, or polysome-high fractions were pooled prior to analysis. Inter-cluster comparisons used a normalized Jaccard Index, which corrects for set size differences:

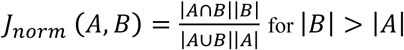

To show that the transcript features associated with translation are independent of the clustering, we identified statistically significantly different transcript isoforms between polysome-low and polysome-high fractions using DESeq2. Polysome-low or polysome-high enriched transcripts exhibited similar trends to those observed when comparing clusters of transcripts (Figure S4B).

### Calculation and comparison of transcript features

Compilation and definition of transcript features was performed as described (Floor and Doudna, 2016). All comparisons are between transcript isoforms derived from the same gene as opposed to all expressed transcript isoforms. Effect size is measured using Cliff’s d, which is related to the Mann-Whitney U statistic by d=(2U/mn)-1, where m and n are the sample sizes of the two sets, and was calculated using the R package orddom (CRAN). Predicted binding sites for brain-specific microRNAs (Shenoy and Blelloch, 2014) were from TargetScan 7.1 (Agarwal et al., 2015).

## Results

### Measuring gene expression during human neuronal differentiation

We differentiated hESCs towards a forebrain fate using a dual-SMAD inhibition protocol (Figure 1A and S1A) (Chambers et al., 2009). We selected four endpoints for analysis: proliferating hESCs, neural progenitor cells (NPCs), cells after 14 days of neural induction, and cells after 50 days of neural induction. These four endpoints were selected to measure initial gene expression in pluripotent stem cells, in proliferating NPCs, in early neural cultures, and more mature cultures containing synaptically active neurons. We collected samples from each endpoint and performed cytoplasmic and nuclear total RNA-seq to measure the transcriptome, ribosome profiling to measure gene-level translation and the locations of ribosomes, and TrIP-seq to measure transcript-specific translation (Table S1). We use the term gene-level translation to indicate the aggregate translation level of all RNA molecules produced from a gene; in contrast, transcript-specific translation refers to an individual RNA transcript produced from a gene.

We confirmed at the mRNA and protein levels that hESCs express the pluripotency marker OCT4 (POU5F1) (Figure 1B,C). NPCs express known markers such as PAX6, SOX1, HES1 and NES (Nestin; Figure 1B,C and S1B), day 14 neural cultures begin to express neuronal genes such as MAP2 (Figure 1B,C and S1B),and more mature (day 50) neural cultures express neuronal markers (CAMKIIA, GRIA1, STMN2, SNAP25, DCX, βIII-tubulin, and synapsin; Figure 1B-D and S1B) as well as glial markers (GFAP, EAAT1 (SLC1A3), and AQP4; Figure S1B). These data reflect the multiple cell types generated during neural induction (Yao et al., 2017). At day 50, we observe strong induction of a class of genes representative of developing human neurons (Pollen et al., 2014); Figure S1C). Therefore, for simplicity, we refer to these as neural cultures. Neurons in 50-day cultures have synaptic puncta, are electrically active, and show excitatory synaptic connectivity (Figure 1D,E).

### Widespread translation changes during neuronal differentiation

To determine the relative contribution of transcription and translation to gene expression changes during neuronal induction, we compared ribosome profiling data to matched total mRNA levels from RNA-seq data. This analysis was performed on hESCs, NPCs, and day 14 neural cultures since it was not possible to obtain ribosome-protected footprints from day 50 neural cultures (see Methods). The ribosome profiling data exhibited the expected preference for protein coding regions, and a peak near the start codon likely caused by translation initiation events trapped by cycloheximide during harvesting of the cells (Figure S2A; (Lareau et al., 2014)). Changes in ribosome profiling and RNA levels were highly correlated between cell types (Figure 2A; r = 0.89 between both hESCs and NPCs or NPCs and day 14 neural cultures), suggesting that transcriptional changes are a major influence on gene-level expression changes during differentiation.

**Figure 2:**
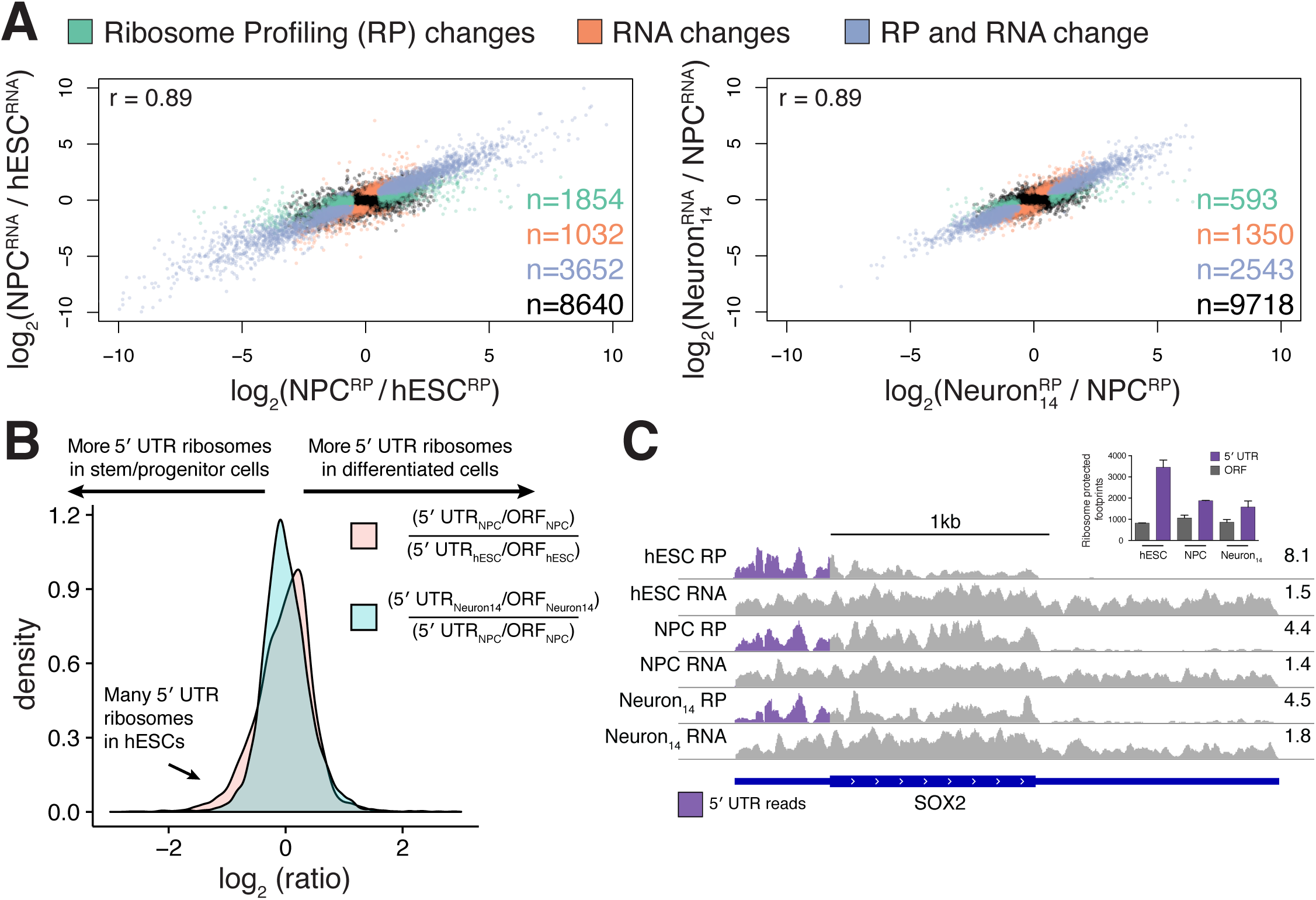
Gene-level translational control during human neuronal differentiation. (A) RNA-seq versus ribosome profiling fold changes between hESCs and NPCs (left) or NPCs and 14-day neural cultures (right) are plotted. Green: differentially expressed (DE) genes in ribosome profiling (RP); orange: DE genes in RNA-seq; blue: DE genes in both RNA-seq and RP; black: genes that do not change expression (p > 0.01). (B) Log-ratios of 5′ UTR to ORF ribosome profiling reads shows high 5′ UTR ribosome density for a group of genes in hESCs compared to differentiated cell types. (C) Relative upstream translation decreasing in the SOX2 gene. Purple: ribosome profiling reads mapping to the 5′ UTR. Right: reads per million. Inset: counts of the ribosome protected footprints mapping to the 5′ UTR or main ORF in different cell types; error is standard deviation. See also Figure S2.

However, thousands of genes exhibited significant celltype-specific changes in translation without corresponding changes in RNA levels, in addition to many genes that changed both transcriptionally and translationally (Figure 2A and Table S2). Analysis of the classes of genes that were differentially regulated at the translational level identified 46 transcription-related genes that were significantly translationally upregulated in NPCs relative to hESCs (e.g. YAF2, ARID1A, CTNNB1 and RELA; Figure S2B), while genes associated with translation itself such as EIF4B, six members of the eIF3 translation initiation factor complex, and 56 ribosomal proteins were significantly translationally downregulated in NPCs (Table S3). Between NPCs and day 14 neural cultures, 593 genes were significantly changed exclusively at the translation level. The most translationally activated genes in neural cultures included the fibronectin FSD1L, the transcriptional repressor JAZF1, and the calmodulindependent protein kinase CAMK4, although no gene ontology terms were significantly enriched (Figure S2B).

Ribosomes can initiate translation at upstream start codons in mRNA 5′ UTRs (uORFs), which affects translation from the major open reading frame (ORF; (Brar et al., 2012; Hinnebusch et al., 2016). We hypothesized that hESC differentiation into neural precursors would result in a redistribution of ribosomes away from mRNA 5′ UTRs, as was observed in mouse embryoid body formation (Ingolia et al., 2011). Indeed, the relative 5′ UTR ribosome occupancy of 148 genes changed by more than a factor of two between hESCs and NPCs (Figure 2B). The distribution of changes was asymmetric, with 101 genes decreasing in 5′ UTR ribosome density in NPCs and only 47 increasing. As an example of this trend, the SOX2 gene, which encodes a SRY-box transcription factor involved in neocortex development (Bani-Yaghoub et al., 2006), has more ribosomes in its 5′ UTR versus the main ORF in hESCs than differentiated cells (Figure 2C). Differentiation of NPCs to neural cultures caused fewer changes in translation within 5′ UTRs, and in the opposite direction to what is found from hESCs to NPCs (82 genes: 54 up in the 5′ UTR and 28 down; Figure 2B). Translation of uORFs can positively or negatively affect translation of the downstream ORF (Brar et al., 2012; Hinnebusch et al., 2016). We found that genes that experienced increased 5′ UTR ribosomes during differentiation generally had decreased ribosome profiling signal in the main ORF and vice versa (Figure S2C). This suggests assembly of translating ribosomes in the 5′ UTR predominantly downregulates translation in this system. Taken together, our results show that transcription is responsible for the majority of gene expression changes during neuronal differentiation while translational control preferentially affects a subset of genes, partly due to relocation of ribosomes from 5′ UTRs in hESCs to open reading frames in differentiated cells.

### Elevated translation of translation-related genes in embryonic stem cells

We identified 172 genes related to translation that were differentially translated between hESCs and NPCs (Figure 3A). mTORC1 and MYC are two major regulators of protein synthesis, so we tested whether they contributed to the enhanced translation of translation factors in hESCs (Bhat et al., 2015; Pourdehnad et al., 2013). MYC levels were elevated in hESCs compared to differentiated cell types (Figure 3B). However, MYC affects transcription of ribosomal protein genes and ribosomal RNA (Ruggero, 2009; van Riggelen et al., 2010) and ribosomal protein genes exhibited relatively small transcriptional changes in these data (Figure 3A). In contrast, mTORC1 post-transcriptionally regulates the translation of a subset of genes that is enriched for translation factors (Thoreen et al., 2012). We therefore assessed mTORC1 activity through phosphorylation of its primary targets involved in translational control, p70S6 kinase and 4E-BP1. We also assessed the phosphorylation state of the p70S6K target, ribosomal protein S6, a commonly used read-out of mTORC1 activity. Differentiation of hESCs into NPCs strongly reduced mTORC1 activity, indicating that suppression of mTORC1 signaling may underlie the downregulation of translational machinery during neural induction (Figure 3C). In support of this, the genes that exclusively experienced translational changes between hESCs and NPCs significantly overlap with those that were previously identified as mTORC1 targets (Thoreen et al., 2012) p = 1.6e-31, hypergeometric test). To experimentally test the role of mTORC1 signaling, we measured the protein levels of genes with elevated translation in Figure 3A after inhibition of mTOR with rapamycin in hESCs (Figure 3D). Protein levels for EIF4B, EIF3D, RPL10A and RPLP1 decrease in abundance following mTOR inhibition, but actin was not affected. Levels of the mTORC1 inhibitor TSC2 increase during differentiation, which is a possible mechanism of the reduction in mTORC1 activity (Figure 3C). We conclude that mTORC1 signaling drives high-level translation of translation-related genes in hESCs.

**Figure 3:**
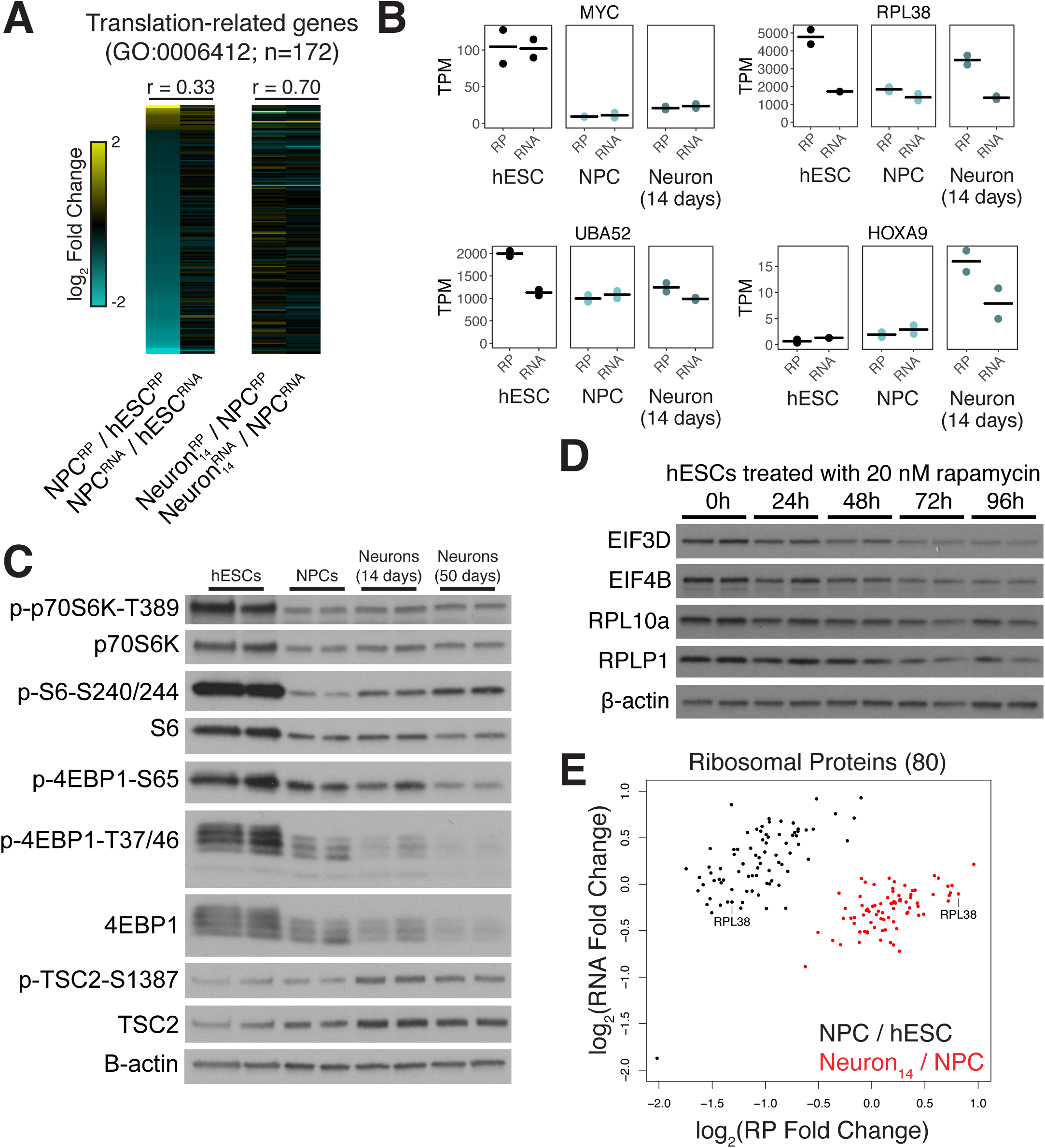
Elevated translation through mTORC1 in hESCs. (A) A heatmap of the log2 fold changes between cell types indicated in either RP or RNA-seq. Cyan: higher in early cell-type; yellow: higher in late cell type. Above: pearson correlation of RP and RNA-seq fold changes. (B) Ribosome profiling and RNAseq expression profiles for MYC, RPL38, UBA52 and HOXA9 across differentiation. Bar: mean expression; points: expression in each replicate. (C) Western blots for phosphop70S6K, phospho-S6, and phospho-4EBP1 as a readout of mTOR activity in differentiating neuronal cells, and the mTOR repressor TSC2. (D) Protein levels from hESCs treated with 20 nM rapamycin for the indicated times in hours for four genes with elevated translation in hESCs in (A). (E) Log2 fold changes in RNA vs ribosome profiling levels for 80 ribosomal proteins between cell types.

We observed that mTORC1 activity increases during early neuronal differentiation (from the NPC state to day 14 neural cultures, Figure 3C). To test whether the reactivation of mTORC1 signaling in early neural cultures correlated with changes in the translational machinery, we compared the translational regulation of ribosomal proteins across hESCs, NPCs, and early neural cultures. Consistent with a reactivation of mTORC1 signaling, translation of many ribosomal proteins increased in early neural cells compared to NPCs (Figure 3B,E). Transcriptional changes were smaller than translational changes for this class of genes between hESCs, NPCs and early neural cells (Figure 3E), highlighting the relative importance of translational regulation in this context.

Active ribosomes can assemble without some ribosomal proteins, leading to the concept of ribosome specialization (Komili et al., 2007; Shi and Barna, 2015). In line with this idea, we found that not all ribosomal proteins change translation level to the same degree between cell types (Figure 3E). For example, RPL38 is strongly translationally upregulated in hESCs versus NPCs, and reactivated in early neural culture (Figure 3B,E). RPL38 selectively controls translation of a subset of genes, including HOX genes (Xue et al., 2015). Intriguingly, translation of the RPL38−-sensitive gene HOXA9 is upregulated in early neural culture when its transcription increases (Figure 3B). Additionally, the fusion gene UBA52, which expresses ubiquitin and RPL40, is translationally upregulated in hESCs compared to NPCs, and RPL40 has also been shown to mediate transcript-specific translation (Figure 3B; (Lee et al., 2013). In sum, our results indicate that mTORC1 signaling drives dynamic ribosomal protein expression during neuronal differentiation, which may in turn yield selective translation of mRNAs in different cellular contexts.

### Transcript-specific translation during human neuronal differentiation

Changes in the translation level of a gene can be driven by altered translation of mRNAs or altered transcript processing, yielding new mRNA transcript isoforms that confer different levels of translation. However, transcript-specific translation is difficult to resolve using ribosome profiling due to the short length of protected footprints, and because alternative 3′ UTRs are invisible to the technique (Ingolia, 2014). To determine the impact of differential transcript isoform expression including 3′ UTRs during human neuronal differentiation, we measured transcript isoform-specific translation using TrIP-seq (Floor and Doudna, 2016) in identical samples to those used for ribosome profiling. TrIP-seq uses polysome profiling and high-throughput sequencing to identify transcript isoforms that are highly translated (polysome-high fraction) and those that are less frequently translated (polysome-low or monosome fractions). Here, fractions were collected from polysome profiles corresponding to the 80S monosome peak, polysome-low (2-4 ribosomes) and polysome-high fractions (5+ ribosomes; Figure 4A). These fractions were chosen because monosome-bound mRNAs are enriched for retained intron and nonsense mediated decay (NMD)-targets compared to polysome-bound mRNAs (Floor and Doudna, 2016; Heyer and Moore, 2016), which represent crucial layers of post-transcriptional control (Lykke-Andersen and Jensen, 2015; Raj and Blencowe, 2015). Furthermore, previous work in human cells demonstrated that polysomal samples clustered into three groups corresponding to the monosome, polysome-low and polysome-high fractions used here (Floor and Doudna, 2016). RNA-seq libraries were made from these fractions and deep sequenced, yielding over two billion mapped reads (Tables S1 and S4).

**Figure 4:**
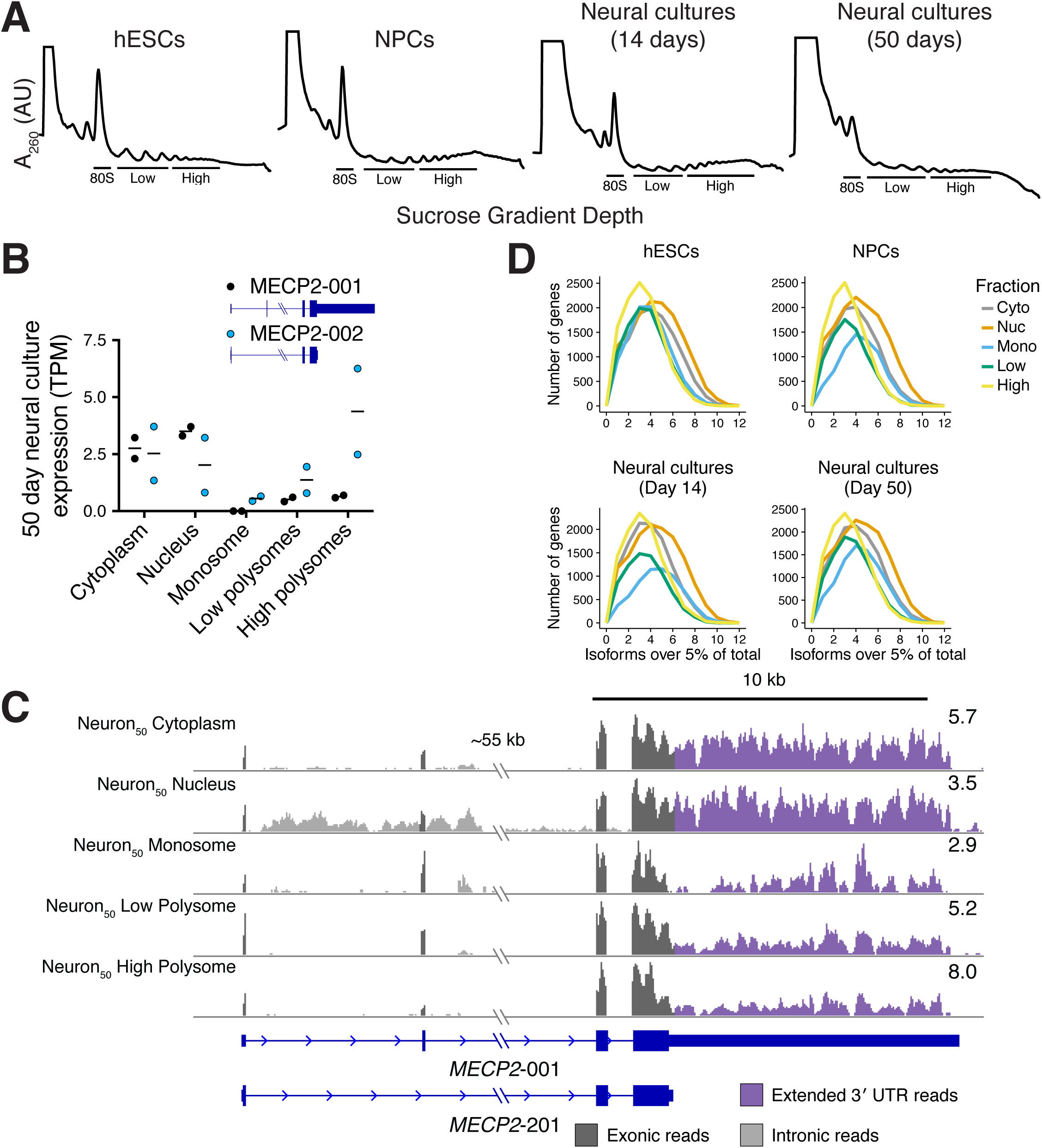
Transcript-level translational control during human neuronal differentiation. (A) Polysome profiles for hESCs, NPCs, day 14 neural cultures and day 50 neural cultures. RNA was collected from the indicated fractions for TrIP-seq. (B) Two differentially translated transcripts of the *MECP2* gene are expressed in 50-day neural cultures, which differ in the length of their UTRs and translation level. Bar: mean expression; points: expression in each replicate. (C) Reads from the *MECP2* locus. Note preferential association of isoforms with the long 3′ UTR and the second alternatively spliced exon with the monosome and low polysome fractions. (D) The number of transcript isoforms per gene expressed above 5% of the total gene expression are plotted in different subcellular fractions.

The striking impact alternative transcript processing has on protein production is evident for genes such as MECP2, which encodes a methyl-CpG DNA binding protein that is altered in a majority of Rett syndrome patients (Lombardi et al., 2015). Some isoforms (e.g. MECP2-001) include an alternative exon that shifts the reading frame of the first start codon in MECP2-002. This exon inclusion therefore introduces a uORF into the mRNA, which downregulates protein production (Figure 4B, (Kriaucionis and Bird, 2004). The 3′ UTR of MECP2 is also alternatively processed, leading to various short (~100 nt) or long (~8 kb) forms, which are differentially translated and differentially expressed during neuronal differentiation (Rodrigues et al., 2016). We observed the impact of both of these transcript processing events in 50-day old human neural cultures. Specifically, reads mapping to the second, alternative exon and the 3′ UTR were higher in the monosome and polysome-low fractions than in the polysome-high fraction (Figure 4C), suggesting lower protein production from these transcript isoforms.

The above example focuses on two transcripts generated by MECP2, but there are 26 annotated isoforms for this gene. In our data, the other transcript isoforms were either not expressed or not found in polysomes, suggesting only a subset of transcript isoforms are translated. We therefore tested whether translation of transcript isoforms is generally pervasive or selective. We computed the number of transcript isoforms that are expressed at more than 5% the total gene expression level in nuclear RNA, cytoplasmic RNA, and the three polysome-derived fractions. We found that nuclear export and translation both act as bottlenecks, selecting a subset of all expressed transcript isoforms. In all four cell stages we found that the nucleus had the highest diversity of transcript isoforms (Figure 4D). A subset of exported transcripts were associated with ribosomes, resulting in diminishing transcripts per gene in cytoplasmic RNA, monosome-associated RNA, and further reductions in transcript diversity in polysomal RNA (Figure 4D). In sum, we find that individual transcript isoforms can vary widely in their translational potential, from being enriched in the nucleus to being highly polysome-associated, and that this variability affects protein production of genes linked to human neurodevelopmental disorders.

### Changes to transcript RNA processing drive gene-level translation changes

To determine the prevalence of transcript-specific translation as in the above MECP2 example, we identified differentially translated transcript isoforms across differentiation. Hierarchical clustering was used to group transcripts by their polysome distribution (Figure 5A, S3B and Table S5). In hESCs and differentiated cell types, there were clusters of transcripts that were primarily associated with either the monosome, polysome-low, or polysome-high fractions. We contrasted these clusters within cell types to identify transcripts of the same gene that were differentially translated. Classifying the types of transcript isoforms in different polysomal clusters identified an enrichment for retained intron transcripts in the monosome fraction in all four cell-types (Figure 5B) (Floor and Doudna, 2016; Heyer and Moore, 2016). We note that the true rate of intron retention is higher as we observed hundreds of unannotated events, for example the cell-type specific intron retention in the DNA methyltransferase DNMT3B (Figure S3A). These retained intron transcripts on polysomes are likely being degraded through NMD as well, which has been suggested to be a mechanism to downregulate classes of genes in neurons (Raj and Blencowe, 2015).

**Figure 5:**
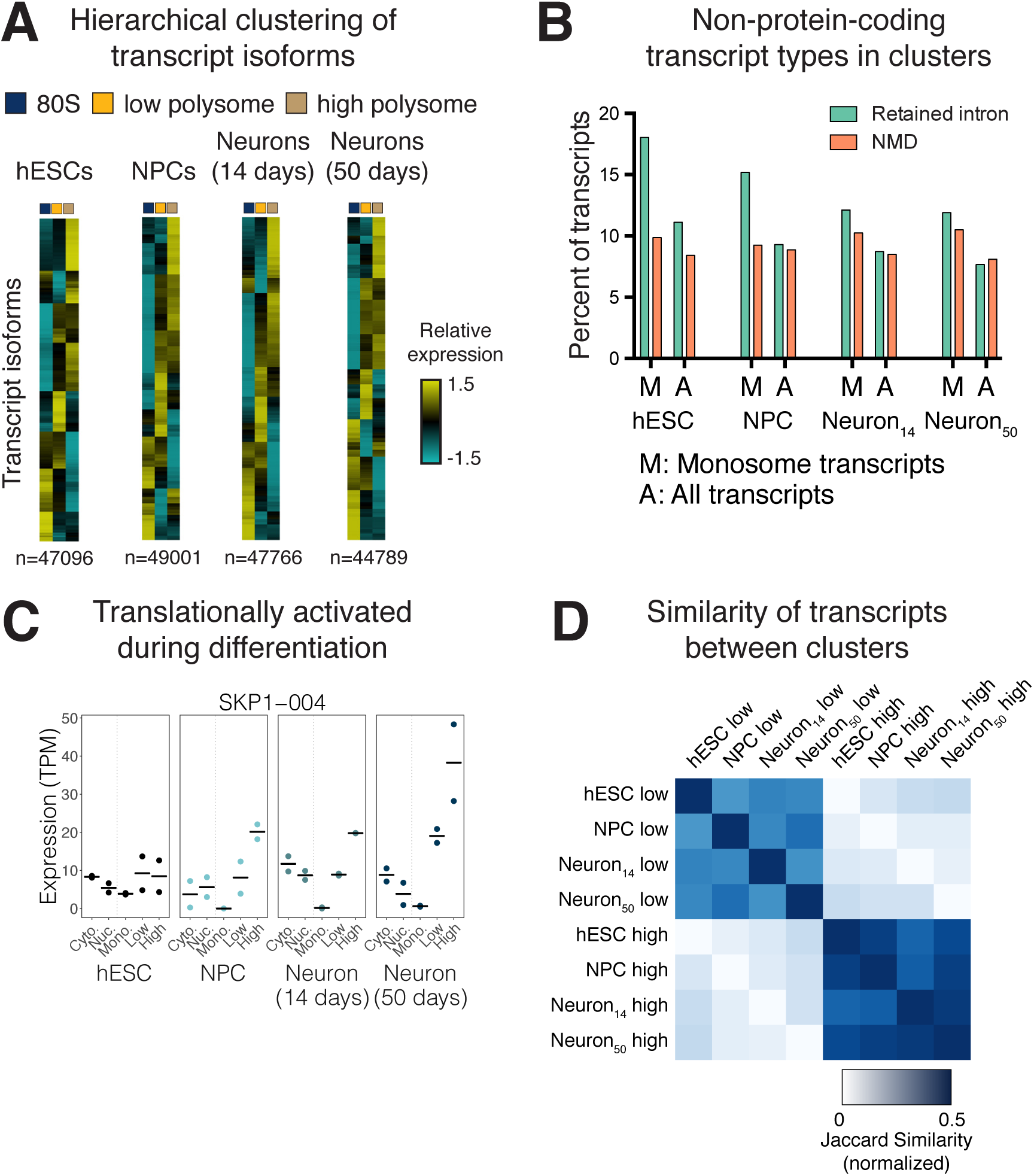
Trends in transcript-level translation during human neuronal differentiation. (A) Heatmaps of hierarchical clusterings of transcript isoform expression. See Figure S3B for dendrograms and average plots. (B) Select transcript types are shown for clusters of transcripts that are primarily present in the monosome fraction. Most other transcripts are annotated as protein coding. (C) Contrasting lowly- and highly-translated clusters between cell types identifies transcripts, such as SKP1-004, that are differentially translated between cell types. Bar: mean expression; points: expression in each replicate. (D) The normalized Jaccard similarity of polysome-low and polysome-high clusters between cell types is plotted. See also Figure S3.

The thousands of translation changes during cell state transitions we observed by ribosome profiling could be driven either by generation of new transcript isoforms with altered translational potency, or altered translation of the same transcript isoforms. To test the relative contribution of these two regulatory mechanisms, we compared clusters of lowly and highly translated transcripts between cell types. Thousands of transcript isoforms moved from low to high (or high to low) polysomal clusters between cell types, likely due to a change in their translation level. For example, a transcript isoform of the SKP1 SCF ubiquitin ligase associated more with the polysome-high fraction following neuronal differentiation (Figure 5C). We analyzed this systematically by computing the Jaccard similarity (the intersection divided by the union) between low and high polysomal clusters between cell types. Between 60-76% of transcript isoforms were detected across pairs of cell types, with day 14 and day 50 neural cultures having the highest similarity. We found that transcript isoforms that were expressed in multiple cell types frequently retained their translation level, even between hESCs and day 50 neural cells (Figure 5D). Thus, changes to steady-state gene-level translation are more frequently driven by production of alternative transcript isoforms rather than changes to the translation level of individual transcripts.

### 3′ untranslated regions repress translation selectively in neural cultures

We next identified features of mRNAs that were associated with differential translation during neuronal differentiation. Even if the coding sequence is identical between two alternatively processed transcripts of a gene, the level of protein output can differ due to changes in the regulatory composition of untranslated regions at the 5′ and 3′ ends (Arribere and Gilbert, 2013; Floor and Doudna, 2016; Hinnebusch et al., 2016; Mayr, 2016; Mayr and Bartel, 2009; Sandberg et al., 2008; Sterne-Weiler et al., 2013). We therefore compared the impact of transcript isoform properties in polysome-low versus polysome-high clusters that might influence translation, such as the density of various regulatory elements in the 3′ UTR or codon usage (Figure S4A). We compared features between isoforms derived from the same gene (Methods). Of the 21 features tested, five affected translation in all cell types, 11 affected translation in a cell type-dependent manner, and five had no or small effects. Long 5′ UTRs were consistently associated with low levels of translation across all cell types (Figure 6A). By contrast, another transcript property, ORF length, was associated with higher ribosome occupancy in all cell types, although this effect decreased in older neural cultures (Figure 6A).

**Figure 6:**
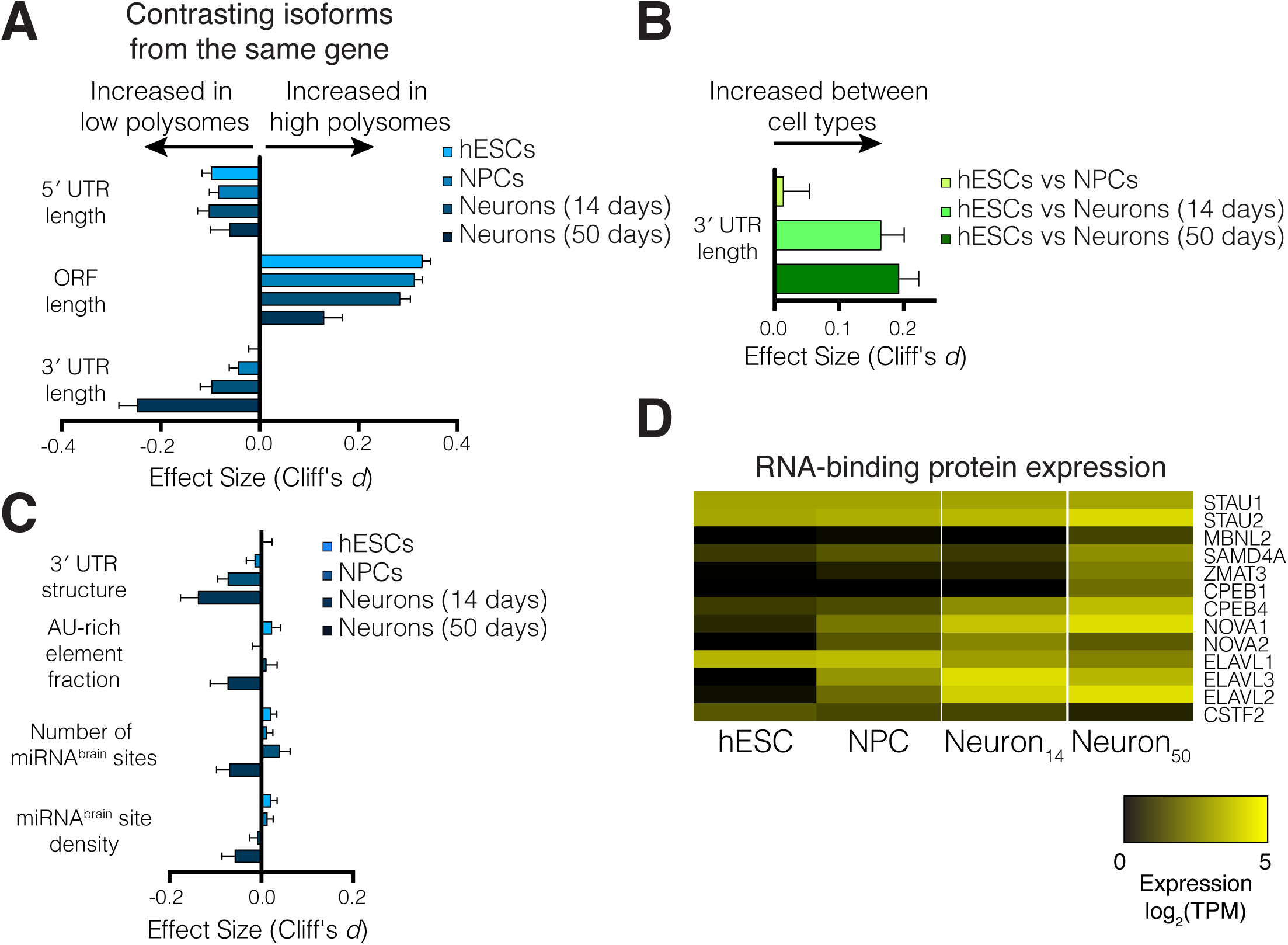
3′ untranslated regions drive differential translation between cell types. (A) The relative length of 3′ UTRs between lowly- and highly-translated transcripts from the same gene increases during differentiation. Error bars: 95% confidence interval, also in (B,C). (B) Relative 3′ UTR length increases for transcripts of the same gene expressed between cell types. (C) 3′ UTR structure, the fraction of 3′ UTRs containing AU-rich elements, and the influence of brain-specific miRNA binding sites increase in 50-day old neurons, suggesting these features may drive translational repression by 3′ UTRs. (D) Expression changes in select RNA binding proteins that influence either 3′ end selection or post-transcriptional control. See also Figure S4.

Transcript 3′ UTR length affected translation more than any other transcript feature tested in 50-day neural cultures (Figure 6A). By contrast, the impact of transcript 3′ UTRs was negligible in hESCs (Figure 6A). Long transcript 3′ UTRs were associated with lower levels of translation as cells differentiated into neural cultures. The simplest explanation for an increased impact of 3′ UTR length on translation during differentiation is changes in 3′ UTR length itself. Alternatively, the effect could be caused by a change in 3′ UTR regulatory content or cellular post-transcriptional control factors. The relative length of 3′ UTRs between transcripts of a gene increased in day 14 neural cultures to a similar extent as day 50 (Figure 6B), yet 3′ UTR length inhibited translation more strongly in day 50 neural cultures than in day 14 cultures (Figure 6A). This suggests changes to the regulatory content of 3′ UTRs or cellular post-transcriptional control factors are responsible for increased translational repression. Indeed, the influence of predicted 3′ UTR secondary structure, the fraction of the 3′ UTR made up of AU-rich elements, and the number and density of binding sites of brain-specific miRNAs (miR-9, -26, - 124, -137, -195, -219-, -338, and let-7; (Shenoy and Blelloch, 2014) all increased during neuronal differentiation (Figure 6C). Furthermore, the expression level of several RNA binding proteins that interact with transcript 3′ UTRs increased during differentiation, such as SMAUG1 (SAMD4A), Staufen1 (STAU1), Staufen2 (STAU2), WIG1 (ZMAT3), and the cytoplasmic polyadenylation element binding proteins CPEB1 and CPEB4. We also saw evidence for changes in gene expression for proteins that regulate alternative polyadenylation, such as CSTF2, multiple ELAV genes, and NOVA1 and NOVA2 (Figure 6D). Taken together, these data indicate that extension of 3′ UTRs during neuronal differentiation induces strong translational repression, which is driven by regulatory elements.

## Discussion

In this work, we analyzed the relationships between transcriptional and translational changes during human forebrain neuronal differentiation. We identified elevated translation of ribosomal protein genes and related translation factors in hESCs, which is driven by mTORC1 signaling. While transcript 3′ UTRs lengthen in early neural cultures, translational repression due to long 3′ UTRs continues to increase as the neural cultures become synaptically active. This is in contrast to long transcript 5′ UTRs, which are generally repressive across all developmental stages. Thus, our work assigns functional consequences to the 3′ UTR extensions that have been observed during brain development, uncovers how post-transcriptional control changes during human neuronal differentiation, and provides a rich resource of untranslated regions that confer global or cell-type-specific protein expression.

Finely tuned translation and mTORC1 signaling are essential for normal brain development, as mTORC1 signaling determines the balance between self-renewal and differentiation of neural stem cells in mice (Hartman et al., 2013; Magri et al., 2011). We find that high mTORC1 signaling in hESCs drives high-level translation of translation-related genes (Figure 3), which decreases during neuronal differentiation. However, hESCs and other stem cell types maintain low overall levels of translation (Fortier et al., 2015; Oh et al., 2005; Sampath et al., 2008; Sanchez et al., 2016; Signer et al., 2014). How might hESCs achieve selective translation of ribosomal proteins and translation-related genes (Figure 3) despite low overall translation? One possibility is that some hESC ribosomes are maintained in an inactive state and are activated upon differentiation into NPCs, for example by dissociation of ribosome assembly factors (Strunk et al., 2012). It is also possible that hESCs preferentially translate a subset of messages through ribosome specialization (Komili et al., 2007; Shi and Barna, 2015; Xue et al., 2015), transcript-specific translation pathways using eIF3 (Lee et al., 2015), or other mechanisms. An interesting future direction will be to define the molecular mechanisms underlying the preferential translation of translation-related genes in hESCs, and to determine its impact on human neuronal differentiation.

Neurons increase their post-transcriptional control potential by extending the set of transcript 3′ UTRs by millions of nucleotides (Hilgers et al., 2011; Miura et al., 2013), but the impact of this expansion on protein production was unknown. Our data clearly indicate that extended 3′ UTRs contribute to translational repression upon neuronal differentiation. Dysregulation of RNA and protein expression in neurons has deleterious consequences and contributes to numerous neurological and psychiatric diseases, affecting all stages of life (Darnell, 2013; Pilaz and Silver, 2015). For example, expression of variants of the TLS/*FUS* RNA binding protein found in amyotrophic lateral sclerosis patients causes aberrant gene expression in human cells, including of *MECP2* (Coady and Manley, 2015). Similarly, copy-number variations in the *NUDT21* gene found in patients with neurological disease increase the fraction of *MECP2* mRNA molecules with long 3′ UTRs (Gennarino et al., 2015). Total *MECP2* mRNA levels are elevated in patient-derived cells (Gennarino et al., 2015) or human cells expressing TLS/FUS variants (Coady and Manley, 2015), but MeCP2 protein levels are decreased in both cases. Thus, measuring translation changes in the manner described here that can detect and quantify alternative transcripts and 3′ UTRs is essential when studying diseases involving genetic alterations to post-transcriptional control factors.

Neurons undergo various forms of plasticity that allow the nervous system to learn and adapt, which often require gene expression changes at both transcriptional and translational levels (Costa-Mattioli et al., 2009; West and Greenberg, 2011). In some cases, local translation at synapses modulates synaptic plasticity (Holt and Schuman, 2013), which involves reactivation of stalled polysomes (Graber et al., 2013). Most methods to measure global translation specifically measure the occupancy of assembled 80S ribosomes on mRNA. Translation is frequently regulated at the initiation step (Hinnebusch et al., 2016), and ribosome occupancy correlates with protein levels (Floor and Doudna, 2016; Ingolia et al., 2009). However, regulated translation elongation decouples ribosome occupancy and protein synthesis, because stalled ribosomes are not actively synthesizing protein. Interestingly, we find that the influence of ORF length on the ribosome occupancy of an mRNA decreases during neuronal differentiation (Figure 6A), which is possibly a sign of increased stalling during elongation. Neuronal elongation stalling is mediated, at least in part, by FMRP binding to the ribosome (Chen et al., 2014; Darnell et al., 2011; Richter and Coller, 2015). Our observation that long transcript 3′ UTRs are enriched in low polysomal fractions (Figure 6) suggests that if these transcripts are incorporated into polysomes that are stalled during elongation, there are few ribosomes bound. Separating the contributions of translation initiation and elongation to neuronal differentiation and function is important, and would be facilitated by approaches that measure the location of multiple ribosomes on a single mRNA.

Our work and previous studies together suggest that celltype-independent control by 5′ UTRs and cell-typespecific control by 3′ UTRs may be a general property of metazoan translation (Floor and Doudna, 2016; Lianoglou et al., 2013; Merritt et al., 2008). However, the regulatory features that control translation both in 5′ and 3′ UTRs are complex (Figure 6 and S4). Our work motivates future experiments measuring translation output from synthetic libraries of untranslated regions (Zhao et al., 2014). Deep sampling of regulatory elements would enable rigorous modeling or machine learning of the dependence of transcript features on translation in different cell types. Such a model would achieve a major goal: to be able to predict the translation level of an mRNA in different cell types based on sequence alone. The complexity of regulatory mechanisms between cell-types identified in this work highlights how important such a model would be, both for understanding basic biology and engineering the translation level of synthetic genes or mRNAs.

Overall, our work reveals extensive changes to gene expression programs at multiple layers during human neuronal differentiation. Our findings facilitate interpretation of alterations to post-transcriptional processes that occur in human neurological and developmental disorders. In summary, multiple aspects of translational control act both in concert with and independently of transcriptional output to shape gene expression during specification of the neuronal lineage.

## Acknowledgements

We thank A. Tambe, B. Do. R.A. Flynn, S. Iwasaki, N. Ingolia, A.S.Y. Lee, S. Venkataramanan, A. Bhate, and S. McDevitt for helpful discussions, sharing reagents, or contributing to software development. We thank B. de Bivort and D. Eckmeier for generating this preprint template. J.D.B. is supported by a Frederick Banting and Charles Best Canada Graduate Scholarship from the Canadian Institutes for Health Research (356733). D.H. is a Pew-Stewart Scholar for Cancer Research supported by the Pew Charitable Trusts and the Alexander and Margaret Stewart Trust. J.A.D. is an HHMI Investigator and a Paul Allen Frontiers in Science investigator. H.S.B. is supported by a fellowship from the Alfred P. Sloan foundation and a NARSAD Young Investigator Grant from the Brain & Behavior Research Foundation. S.N.F. is a HHMI fellow of the Helen Hay Whitney Foundation. This work used the Vincent J. Coates Genomics Sequencing Laboratory at UC Berkeley, supported by the NIH S10 OD018174 Instrumentation Grant. This work was supported by funds to H.S.B. from the Hellman Family Faculty Fund and NIH R01NS097823 and to D.H. through the Siebel Stem Cell Center and NIH R01CA196884.

---

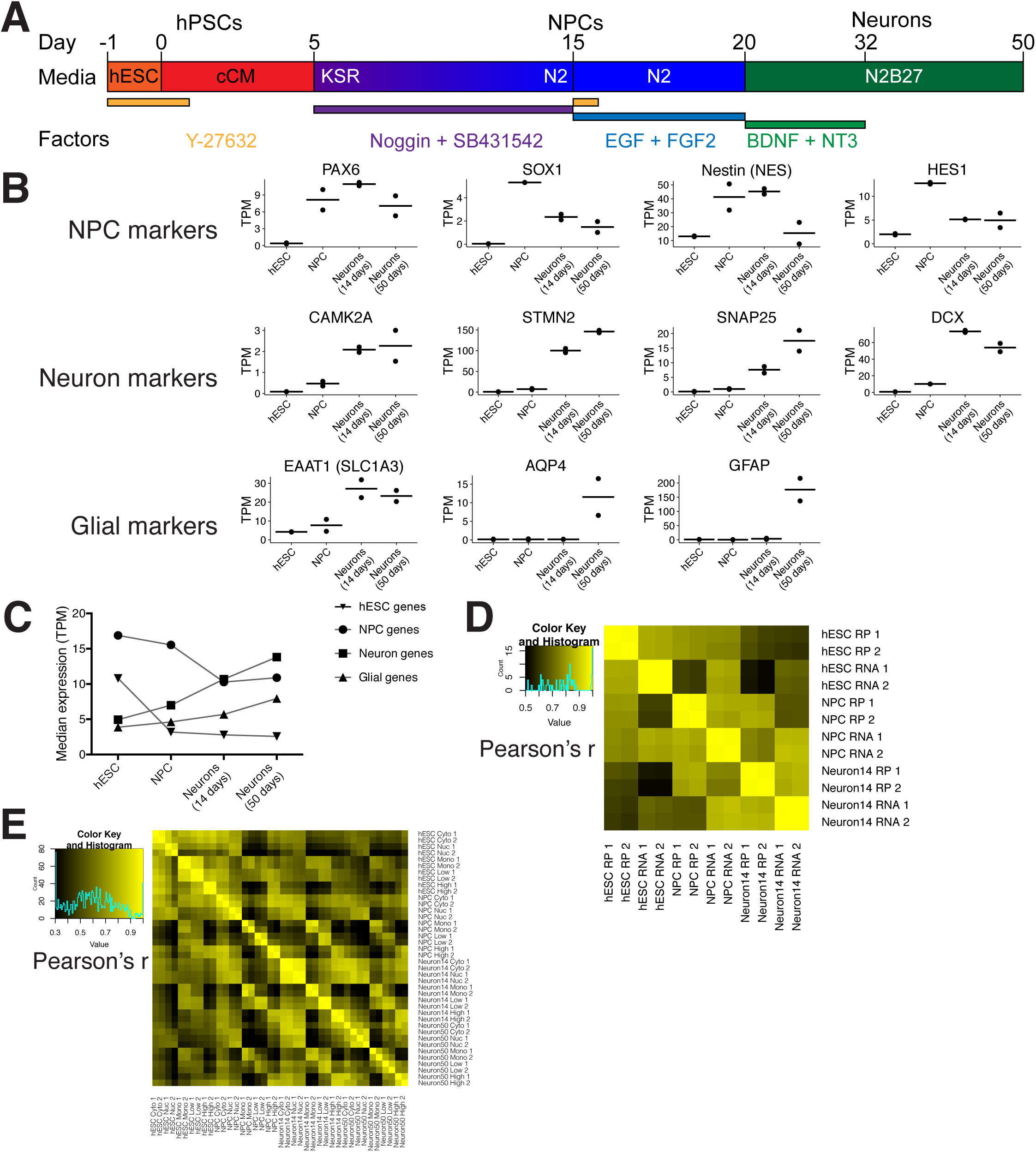
Figure S1, related to Figure 1: (A) A schematic of the neural induction protocol. (B) Marker gene expression in cytoplasmic RNAseq for eleven genes. TPM: transcripts per million. Bar: mean expression; points: expression in each replicate. (C) Median gene expression for classes of genes representing different forebrain cell types. Classes of genes derived from single cell sequencing of human fetal brains in Pollen et al. 2014 or hESC profiling in Mallon et al. 2013. (D) Replicate correlations for ribosome profiling and matched RNAseq samples. (E) Replicate correlations for cytoplasmic, nuclear, and polysomal (TrIP-seq) RNAseq samples.

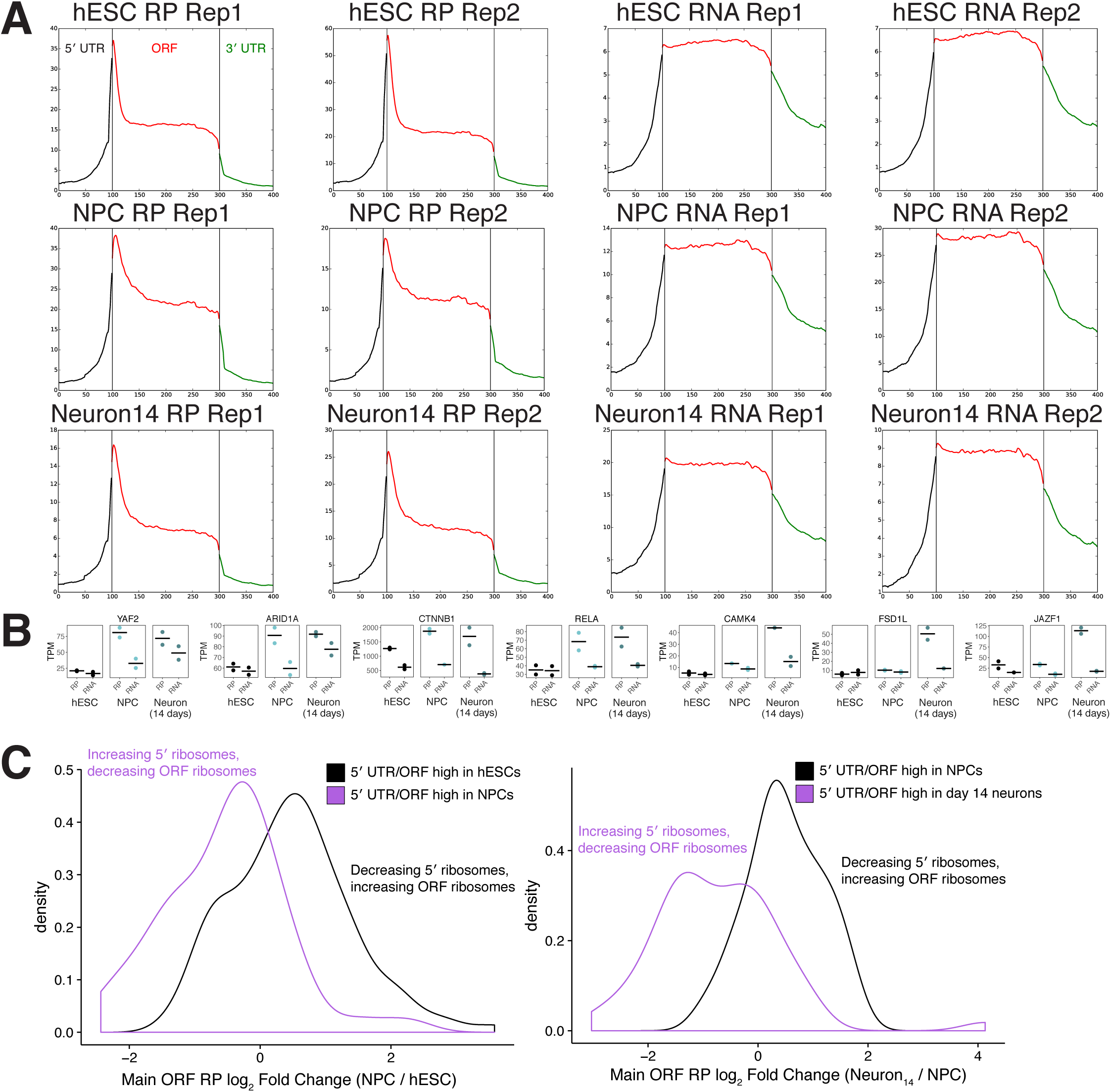
Figure S2, related to Figure 2: (A) Metagene plots for each replicate of ribosome profiling and RNAseq in hESCs, NPCs and 14 day neural cultures. Metagene plots are constructed by collapsing each ORF to 200 bins and averaging RP density across genes, while 5′ and 3′ UTR regions are the 100 nt before and after each ORF averaged across genes. (B) RP and RNAseq gene expression changes for seven genes related to transcription (left four) or translationally activated in day 14 neural cultures (right three). (C) Fold change distributions in ribosome profiling on the main ORF for classes of genes with changes in 5′ UTR /ORF occupancy during differentiation. Note the reciprocal relationship between 5′ UTR ribosomes and ORF ribosomes.

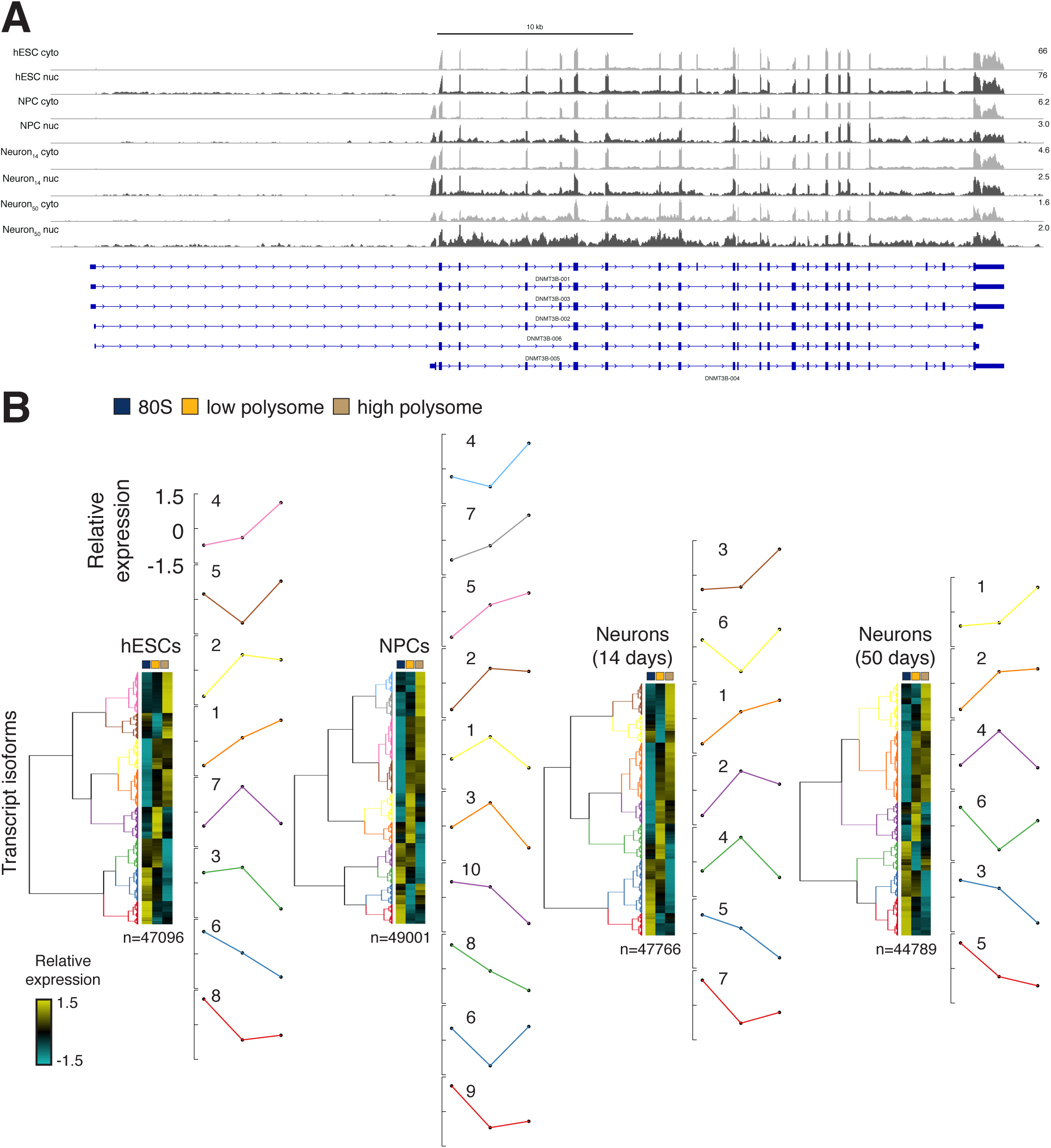
Figure S3, related to Figure 5: (A) Unannotated intron retention increases in DNMT3B during neural induction. Gray: cytoplasmic RNAseq. Dark gray: nuclear RNAseq. Note also the cell type specific exon inclusion in, for example, the 2nd and 3rd to last exons, among others. (B) The clustering in Figure 5A is reproduced here with associated dendrograms and plots of average expression of all transcripts in each color. Colors of the expression profiles match the colors of the dendrogram. All average plots run from 1.5 to -1.5 in units of relative expression (rlog; see Methods). Cluster IDs are the numbers to the upper left of each cluster average plot.

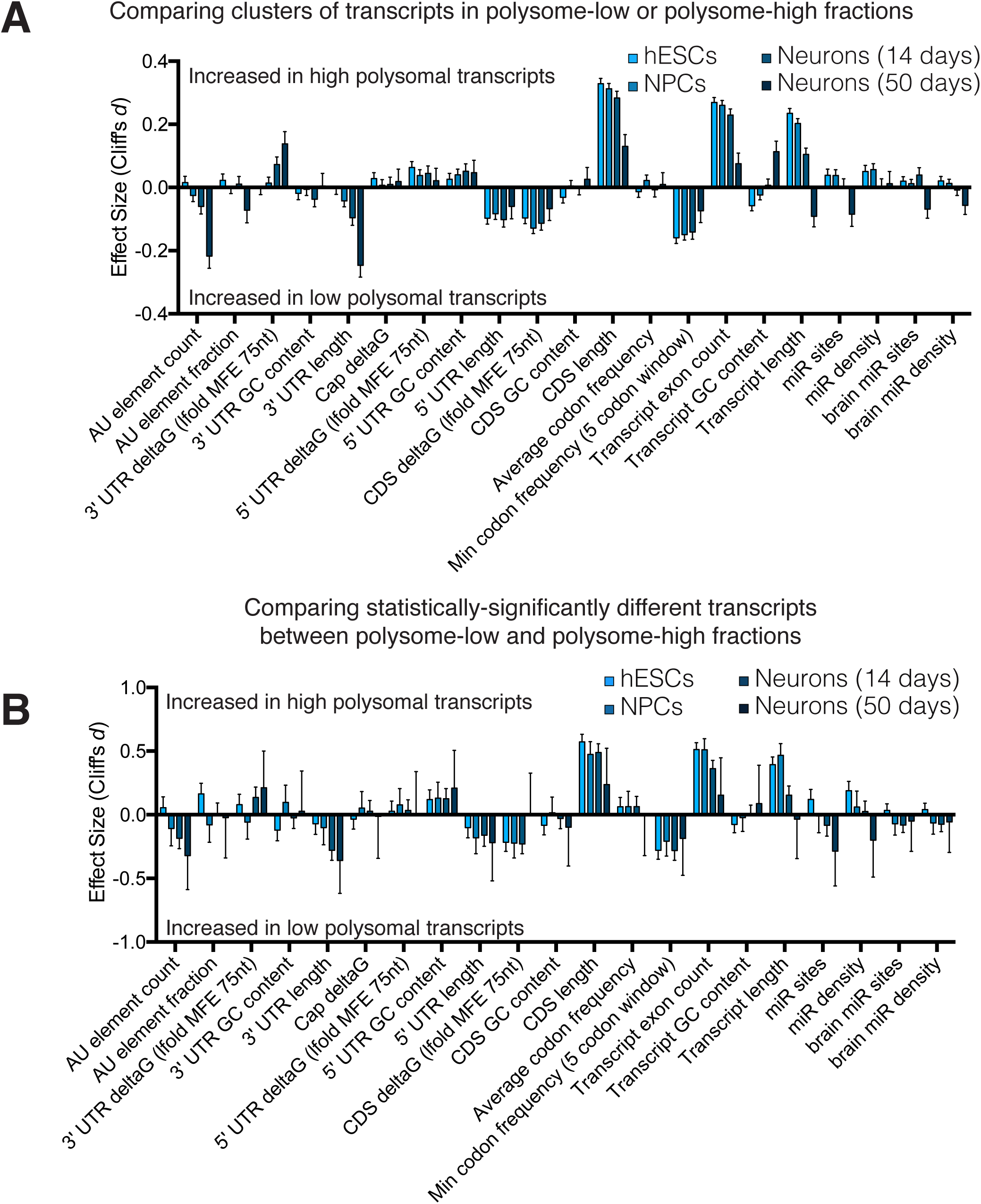
Figure S4, related to Figure 6: (A) The effect size of 21 different features between transcripts in high polysomal clusters versus those in low polysomal clusters. Different transcript isoforms of the same gene were contrasted. Error bars are 95% confidence intervals. MFE 75nt: minimum free energy across a 75 nucleotide sliding window in the region. Codon frequency is the usage of codons in the human transcriptome. Note: higher deltaG for a set of transcripts implies less stable structures. See Methods for more details. (B) as in (A) but for transcripts identified by DESeq2 as significantly different (p < 0.01) between low polysomal fractions and high polysomal fractions.

